# Item-specific memory reactivation during sleep supports memory consolidation in humans

**DOI:** 10.1101/2023.01.25.525599

**Authors:** Jing Liu, Tao Xia, Danni Chen, Ziqing Yao, Minrui Zhu, James W. Antony, Tatia M.C. Lee, Xiaoqing Hu

**Author notes:** Correspondence should be sent to: T.M.C. L.,; X. H.

## Abstract

Memory consolidation stabilizes newly acquired information. Understanding how individual memories are reactivated during sleep is essential in theorizing memory consolidation. Via unobtrusively re-playing auditory memory cues to sleeping human participants, we identified the reactivation of individual memories during slow-wave sleep (SWS). Using representational similarity analysis (RSA) on cue-elicited electroencephalogram (EEG), we found functionally segregated item-specific representations: the early post-cue EEG activity (0-2 seconds) contained comparable representations for memory cues and for non-memory control cues, thus reflecting sensory processing. Critically, the later EEG activity (2.5-3 s) showed greater item-specific representations for post-sleep remembered items than for forgotten and control cues, demonstrating the reactivation and consolidation of individual memories. Moreover, spindles preferentially supported item-specific memory reactivation for items that were not tested before sleep. These findings delineated how cue-triggered item-specific memory reactivation, subserved by spindles during SWS, contributed to memory consolidation. These results will benefit future research aiming to perturb specific memory episodes during sleep.

## Introduction

Newly acquired experiences must undergo consolidation to become long-lasting memories (*1*). Studies using various techniques (e.g., single-unit recordings, fMRI, EEG) suggest repeated, covert reactivation of memories occurs during post-learning non-rapid eye movement (NREM) sleep (*2–8*). Intriguingly, memory reactivation can be manipulated by unobtrusively re-presenting sensory cues associated with wakeful learning during subsequent sleep, a paradigm known as targeted memory reactivation (TMR) (*9–12*). TMR allows researchers to determine when and which memories become reactivated via re-playing stimulus-specific memory cues, which shows promise in modulating the declarative, procedural, and emotional memories, in both clinical and educational settings (*10, 13–16*). However, given the challenges in identifying memory-specific neural ensembles during sleep in humans (*17*), how individual TMR cues reactivate their corresponding memory representations and thereby facilitate memory consolidation remains unclear.

Recent studies using multivariate neural decoding methods have advanced our understanding of TMR-based memory consolidation. First, TMR cue-elicited neural activity contained task-related or category-level memory representations (*18–21*). Second, during NREM sleep, neural representations associated with wakeful retrieval re-emerged rhythmically ∼1-Hz following TMR cues (*22*). While these findings suggest that memory reactivation can be detected during NREM sleep, whether TMR cueing triggers the fined-grained neural representations for individual memories remains unknown. Moreover, although the sleeping brain remains its robust neural auditory responses to external stimuli (*23, 24*), how processing auditory memory cues drives the exogenous memory reactivation during sleep is yet to be established.

Converging evidence suggests that the thalamocortical spindles are instrumental for both endogenous and exogenous memory reactivation during NREM sleep (*25, 26*). In TMR, enhanced post-cue spindle-related sigma power not only predicted TMR effects (*27, 28*) but is also correlated with the distinctiveness of category-level memory representations (*19*). Furthermore, TMR effects are abolished when post-cue sigma power is reduced (*29*) and when cueing occurs during the spindle refractory period, i.e., ∼3-6 s following a spindle when a spindle is less likely to occur (*27*). This evidence prompts an intriguing question of whether the spindle in specific time windows can even support memory reactivation for individual items.

An important factor that may modulate TMR effects and spindle-mediated memory consolidation is pre-sleep testing. Given that pre-sleep testing could enhance memory via retrieval-induced fast consolidation processes (*30, 31*), testing may reduce the subsequent TMR effects (*32*) and weaken the relationship between the neural activity and memory consolidation. Indeed, both spontaneous sleep and sleep TMR preferentially benefit weaker memories, with sleep spindles consolidating weak rather than strong memories (*33–36*). Alternatively, pre-sleep testing may enhance the future relevance of the tested items, which would make these memories preferentially consolidated due to their motivational salience (*37, 38*). To reconcile these competing hypotheses, we further examined how pre-sleep testing affects item-specific memory reactivation and spindles.

We used representational similarity analysis (RSA) on scalp electroencephalogram (EEG) (*19, 39, 40*) to extract item-specific representation during slow-wave sleep (SWS). Employing a subsequent memory analysis approach to link item-specific representations during sleep with post-sleep memory performance, we identified two functionally distinct time windows.

Specifically, the early time window (0-2 s post-cue) exhibited item-specific representations for both the memory cues and the non-memory control cues. However, item-specific representations in this early time window showed no subsequent memory effect or any difference between memory items and control items, thus primarily reflected sensory processing of individual auditory cues. More importantly, a later 2500-2960 ms time window bore relevance with post-sleep memory performance: remembered items exhibited greater item-specific representations than both forgotten and control cues. Notably, sleep spindles during this time window supported item-specific memory reactivation only for pre-sleep untested items, highlighting the nuances of spindle-mediated mechanisms in memory consolidation.

## Results

### Behavioral results

Thirty participants (23 females, mean ± SD age, 22.37 ± 2.94 years) were included in the analyses (see Materials and Methods for participants exclusion criterion). Participants first studied 96 word-picture (cue-target) pairs, with each pair being repeated three times. After a distractor (math) task, participants completed a pre-sleep cued recall test, with half (48) of the learning pairs being tested (i.e., tested items) while the other half (48) were untested (i.e., untested items). TMR cues consisted of 24 spoken words from each of the tested and untested items, respectively, resulting in a total of 48 TMR cues. For tested items, pre-sleep cued recall performance was matched for subsequently cued and uncued items. For untested items, cues were pseudo-randomly selected (see Materials and Methods). In addition to these 48 TMR cues, four spoken words that were not paired with any learning content were included as non-memory control cues, and they were played the same number of times as TMR cues during sleep. On the following morning, memory was tested for all 96 learned pairs (see Fig. 1).

**Fig 1.**
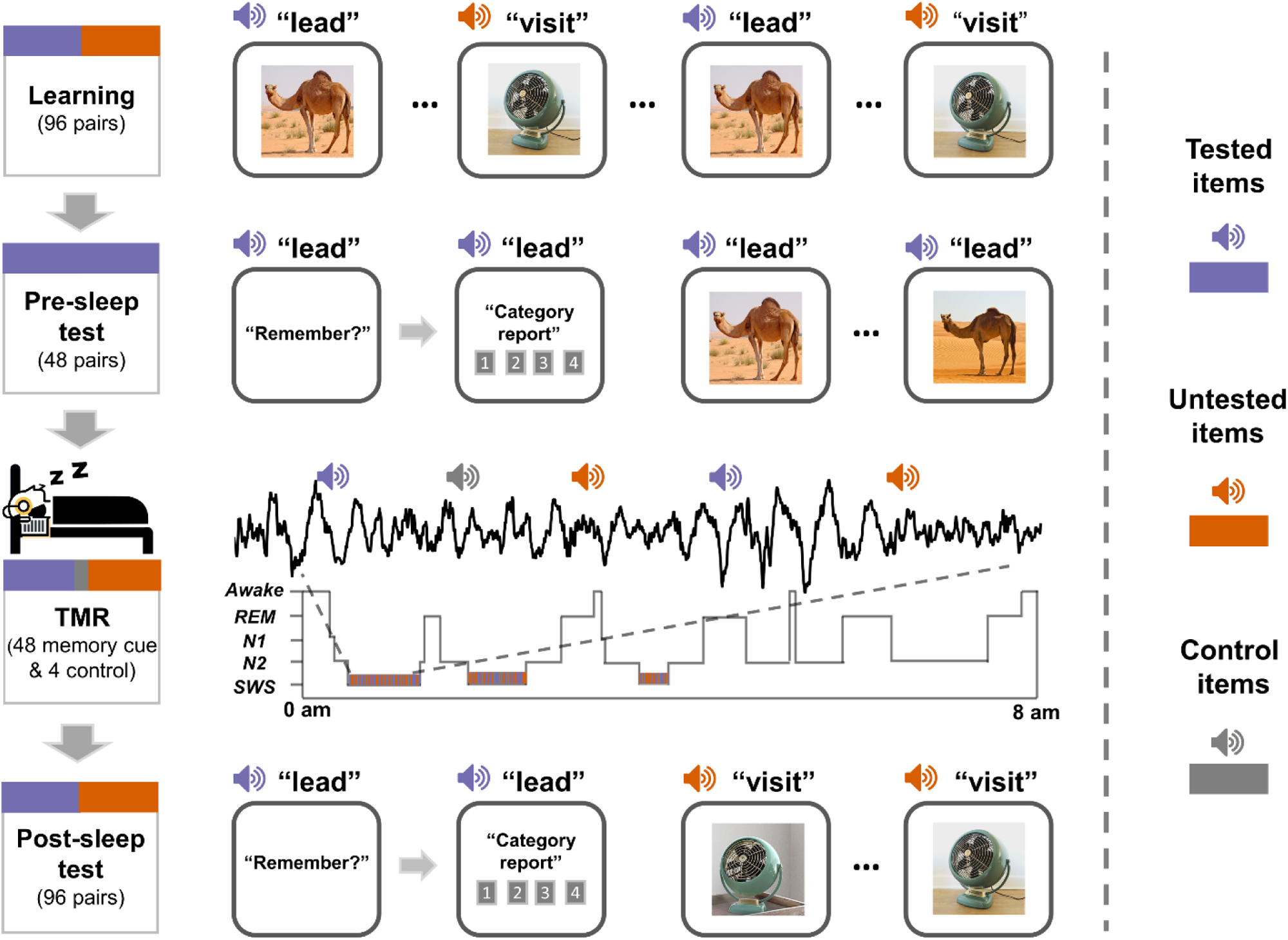
Experimental design. The experiment included four main phases, i.e., learning, a pre-sleep test (cued recall and recognition), targeted memory reactivation (TMR) during slow-wave sleep (SWS), and a post-sleep test. During the learning phase, participants studied word-picture pairs, half of which were tested before sleep (blue rectangle) while half were not (orange rectangle). Auditory cues from half of the tested items and half of the untested items, along with four control cues (grey rectangle), were played during SWS of the overnight sleep. Memory for all pairs was assessed during the post-sleep test (cued recall and recognition) the next morning.

We first examined how pre-sleep testing affected TMR effects by conducting a 2 (cued vs. uncued) by 2 (tested vs. untested) repeated measures ANOVA on post-sleep memory recall performance. Results revealed neither an interaction effect (*F*(1,29) = 0.15, *p* = 0.700) nor the TMR main effect (cued vs. uncued, *F*(1,29) = 0.11, *p* = 0.739) (see Fig. S1). The same ANOVA on the post-sleep memory recognition performance similarly yielded no effects (all *p*s > 0.373, see Fig. S1). Despite the absence of significant TMR effects at the group level, substantial individual differences were observed (see Fig. S1), which prompted further investigations into individual differences analyses of TMR effects. In the following analyses, we used cued-recall performance, which is more sensitive than the recognition performance to TMR effects (*10*).

### Early neural responses to auditory cues predict TMR effects

Previous studies demonstrated that external auditory stimulation during NREM sleep can increase EEG power in theta (∼4-7 Hz), alpha (∼8-12 Hz), and sigma (∼11-16 Hz) bands (*19, 41, 42*). In line with these studies and confirming that the auditory cues were indeed processed during sleep, we found that all memory cues (i.e., including both the tested and untested items) significantly enhanced EEG power in a broad range of 2-40 Hz during the first 1960 ms as compare to pre-cue baseline (*p*_corr_ < 0.001, see Fig. 2A). Notably, two prominent frequency ranges emerged: a low-frequency range of 2-9 Hz and an extended sigma band of 11-18 Hz. Following this early cluster, we observed a significant reduction of sigma power in a late cluster (2320-3380 ms, *p*_corr_ = 0.005), which potentially reflected the spindle refractoriness (*27*). Further tests suggested that pre-sleep testing did not modulate cue-elicited EEG power in the above clusters (all *p*s > 0.102, see Fig. S2). Together, these results suggest that memory cues were indeed processed by the sleeping brain during NREM sleep.

**Fig 2.**
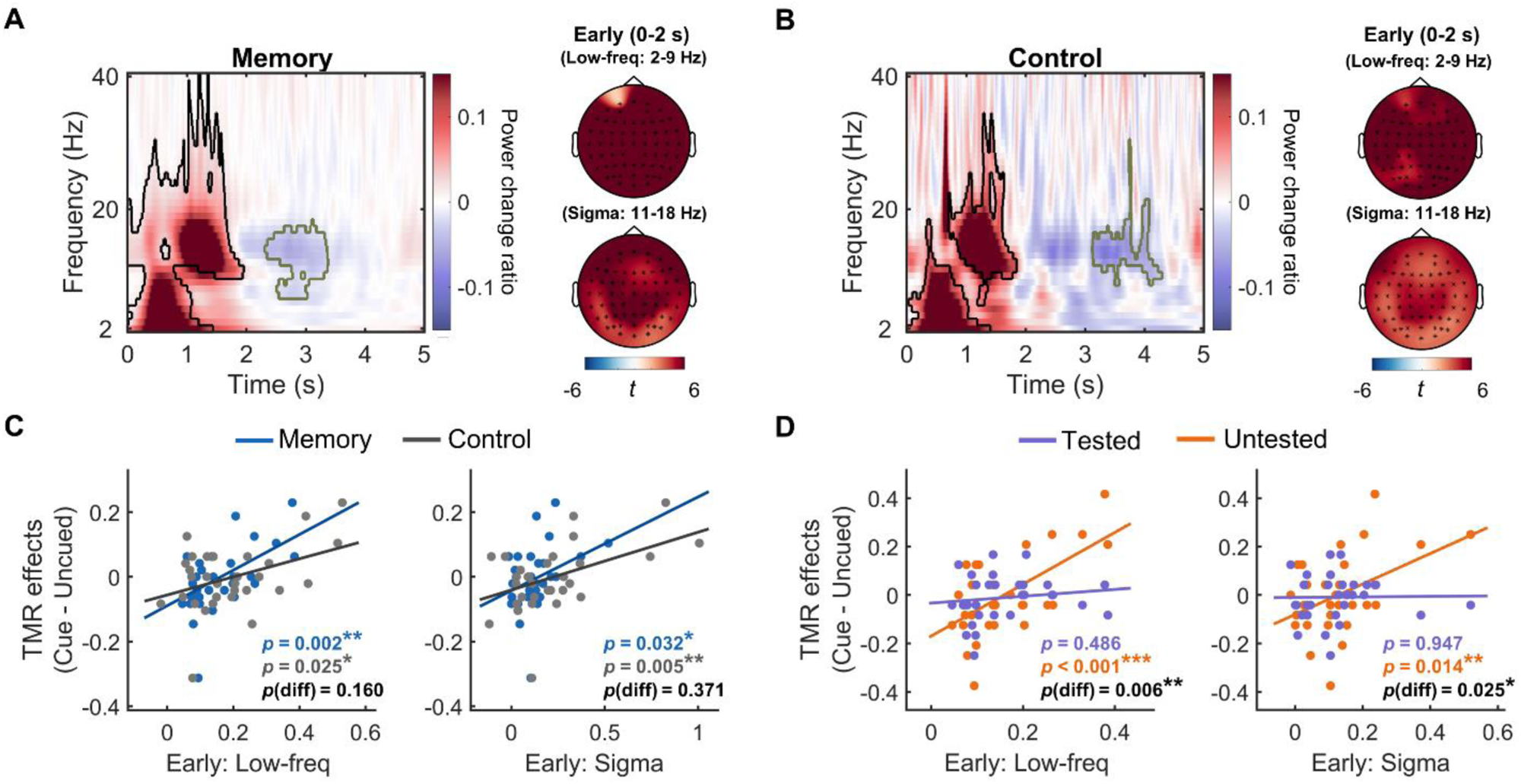
TMR cue-elicited EEG spectral power changes in the early time window (i.e., within the first 2 seconds after cue onset) and their association with post-sleep TMR effects. **(A)** Left panel: memory cues enhanced EEG spectral power across all memory items within an early cluster (circled by the black line), followed by decreased sigma power in a late cluster (circled by the green line), compared to the pre-cue baseline. Right panel: topography plots for the memory cue-elicited low-frequency range and sigma band power in the early cluster. **(B)** Left panel: non-memory control cues enhanced EEG spectral power within the two early clusters (circled by the black line), followed by decreased sigma power in a late cluster (circled by the green line), compared to the pre-cue baseline. Right panel: topography plots for the control cue-elicited low-frequency range and sigma band power in the early clusters. **(C)** Both memory cue and control cue-elicited EEG power in the low-frequency range and the sigma band in the early cluster positively predicted post-sleep TMR effects. The prediction effect did not differ between memory and control cues. **(D)** Memory cue-elicited EEG power in the low-frequency range and the sigma band in the early cluster predicted the post-sleep TMR effects for untested but not for tested items. The *p*-values indicate the significance of robust linear regression that minimized the potential influence of statistical outliers. *: *p* < 0.05; **: *p* < 0.01; ***: *p* < 0.001.

To further investigate whether the enhanced EEG power reflects memory-specific processing or sensory processing to auditory cues, we examined the EEG power locked to the non-memory control cues. Results showed that control cues also enhanced EEG power in the low-frequency range and the sigma band within the first 2 s (both *ps*_corr_ < 0.001), followed by sigma power reduction (3130-4300 ms, *p*_corr_ = 0.006) (Fig. 2B). Importantly, direct comparisons between the EEG power elicited by memory cues and control cues after matching trial numbers revealed no significant differences (*p*_corr_ > 0.441, Fig. S2). These findings indicated that the early neural responses reflected how the sleeping brain reacts to external auditory stimuli, regardless of memory cues or not.

We next examined whether the cue-elicited EEG power could predict the post-sleep TMR effect (i.e., cued minus uncued recall performance). Using robust linear regression analyses that minimized the influence of potential statistical outliers, results revealed that memory cue-elicited EEG power in the low-frequency range (2-9 Hz) and the sigma band (11-18 Hz) in the early cluster (within first 2s) positively predicted TMR effects (early low-frequency: adjusted *R*^2^ = 0.28, *p* = 0.002; early sigma: adjusted *R*^2^ = 0.12, *p* = 0.032, Fig. 2C). More specifically, theta (4-7 Hz), and alpha (8-12 Hz) power significantly predicted TMR effects (all *p*s < 0.026). However, sigma power reduction during the later time window (i.e., 2320-3380 ms) did not predict TMR effects (adjusted *R*^2^ = -0.004, *p* = 0.363). Interestingly, EEG power elicited by non-memory control cues showed similar prediction effects: early, but not late EEG power positively predicted the TMR effects (early low-frequency: adjusted *R*^2^ = 0.14, *p* = 0.025; early sigma: adjusted *R*^2^ = 0.22, *p* = 0.005; late sigma: adjusted *R*^2^ = 0.064, *p* = 0.097, Fig. 2C). Directly comparing the prediction effects of memory and control cues showed no significant differences (all *p*s > 0.159).

Intriguingly, pre-sleep testing remarkably modulated the relationship between cue-elicited EEG power and TMR effects. Specifically, cue-elicited EEG power significantly predicted TMR effects only for untested items (early low-frequency: adjusted *R*^2^ = 0.32, *p* < 0.001; early sigma: adjusted *R*^2^ = 0.17, *p* = 0.014; Fig. 2D), but not for tested items (both *p*s > 0.485). More importantly, this prediction effect was significantly higher for untested items than for tested items (both *p*s < 0.026). Again, EEG power elicited by non-memory control cues showed similar prediction effects for pre-sleep tested and untested items (see Fig. S2). Taken together, these results suggested that a stronger early neural response to external auditory stimuli during NREM sleep predicts TMR efficacy. This association was observed regardless of whether these auditory stimuli were memory cues or non-memory control cues. Furthermore, this association was particularly strong for untested items but not for tested items.

### Early EEG activity contains item-specific representations for cue processing

Demonstrating that this early EEG power was elicited by both memory and non-memory control cues, we hypothesize that this early EEG activity might contain neural representations of individual auditory cues. To test this hypothesis, we performed the representational similarity analysis (RSA) between trials of the same cues (within-item [WI] similarity) as well as between trials of different cues (between-item [BI] similarity) (Fig. 3A). We computed the representational similarity by correlating the cue-elicited raw EEG pattern (filtered between 0.5-40 Hz) across all channels within consecutive overlapping time windows of 500 ms, with a 10 ms sliding step, during a 5-second time window after cue onset. The WI and BI similarity were calculated between trials from two different TMR blocks to control for the temporal proximity effect. Item-specific representations were indicated by significantly greater WI similarity than BI similarity. Our analysis revealed significant item-specific representations during 560-1350 ms after cue onset (*p*_corr_ = 0.008, Fig. 3B). The same analyses on control items similarly revealed significant item-specific representations during 1440-1950 ms after cue onset (*p*_corr_ = 0.040, Fig. 3B). Notably, the relatively delayed onset of item-specific representations for control cues compared to memory cues may reflect the cost of processing speed for the control cues given they were less studied pre-sleep (*43*). Moreover, a repeated measures ANOVA on item type (memory vs. control) and item-specificity (WI vs. BI) did not reveal any significant clusters (*p*_corr_ > 0.085).

**Fig 3.**
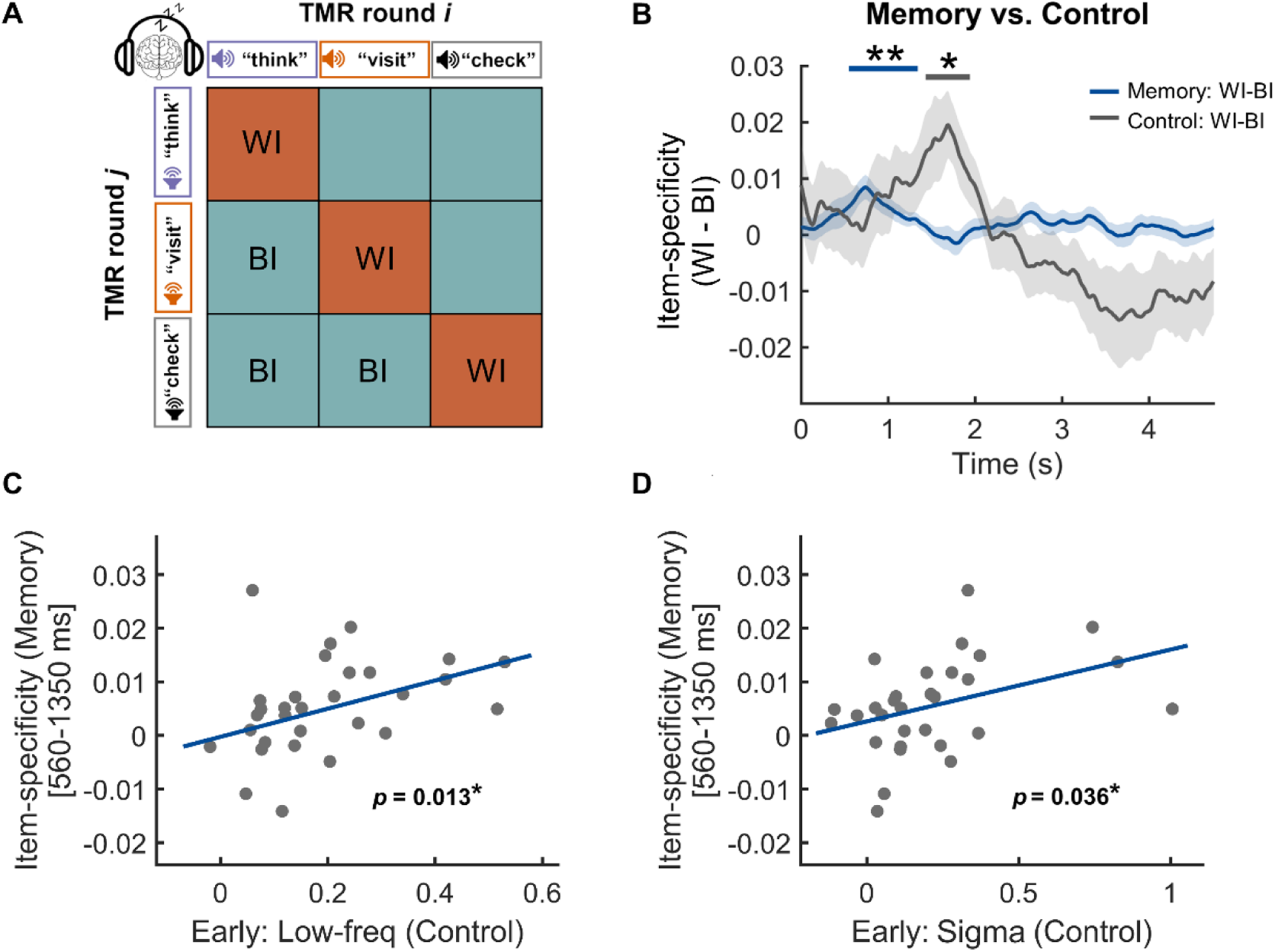
Item-specific representations following TMR cues during slow-wave sleep. **(A)** Representational similarity analysis scheme across different TMR blocks. Artifact-free raw EEG data pattern was correlated between trials with the same auditory cues (within-item similarity, WI) and between trials with different auditory cues (between-item similarity, BI). Item-specific representations were examined by contrasting the WI similarity versus BI similarity. **(B)** Item-specific representations for memory items and control items were significant in the time windows 560-1350 ms (blue horizontal bar on the top) and 1440-1960 ms (gray horizontal bar on the top) after cue onset, respectively. **(C-D)** Control cue-elicited low-frequency EEG power and sigma band EEG power in the early cluster (< 2 s, see Fig. 2B) predicted the item-specific representations for memory items in the 560-1350 ms time window, respectively. *: *p* < 0.05; **: *p* < 0.01.

We next performed the linear regression analysis between control cue-elicited EEG power, which were sensory-cue driven neural responses, and memory cue-elicited item-specific representations. The results revealed that both the control cue-elicited low-frequency range and sigma band EEG power in the early time window significantly and positively predicted the memory cue-elicited item-specific representations in the 560-1350 ms time window (Low-frequency: adjusted *R*^2^ = 0.18, *p* = 0.013; Sigma: adjusted *R*^2^ = 0.12, *p* = 0.036; see Fig. 3C-D). Additional control analyses also found that memory cue-elicited EEG power in an earlier non-overlapped time window positively predicted the following item-specific representations for memory items (*p* < 0.001, see Fig. S3). A similar significant positive prediction effect was also observed for control items (*p* = 0.024, see Fig. S3).

These results provided converging evidence that the early enhanced EEG power within the first 2 s reflected effective processing of individual auditory cues. Furthermore, the control cue-elicited EEG power predicted item-specific representations of memory items, highlighting that the level of the sleeping brain responding to external auditory cues is crucial in shaping item-specific representations of these cues.

### Later EEG activity contain item-specific representation for memory reactivation

We next examined our key question: whether TMR cues reactivate item-specific memory representations during sleep that mediate memory consolidation. Adopting a subsequent memory approach, we conducted a memory (remember vs. forget) by item-specificity (WI vs. BI) repeated measures ANOVA. The results indeed revealed a significant interaction effect during the 2500-2960 ms time window (*p*_corr_ = 0.025, Fig. 4A), suggesting that TMR cues reactivate item-specific memory representations. Notably, the observed item-specific representations were above and beyond category-level representations, given that the WI similarity was greater than the BI similarity even when the BI items were drawn from within-category trial pairs (i.e., within-category similarity, WC, see Fig. S4).

**Fig 4.**
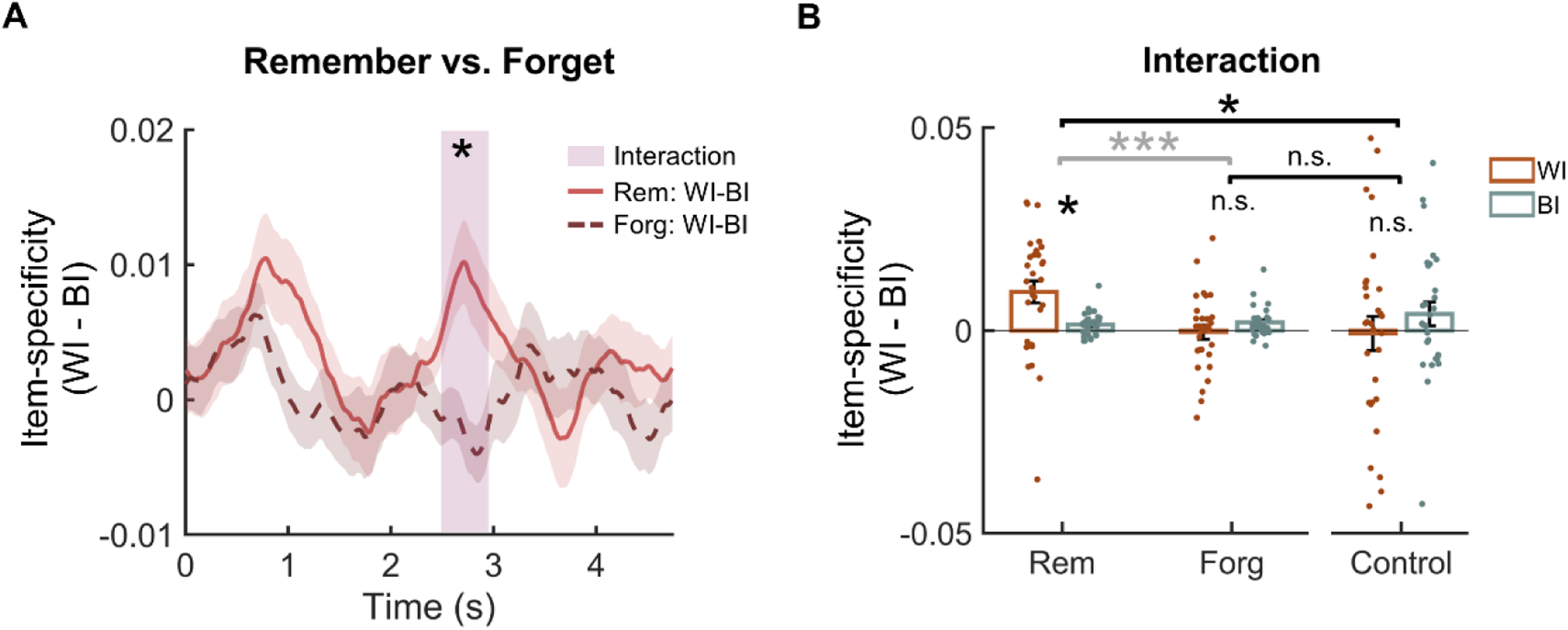
Item-specific memory reactivation. **(A)** Item-specificity by post-sleep memory repeated measures ANOVA revealed a significant interaction effect cluster in 2500-2960 ms time window (shaded rectangle). **(B)** Post-hoc analyses of the interaction cluster (the shaded rectangle in A) found WI similarity significantly greater than the BI similarity for remembered items, while there were no differences between WI and BI similarity for forgotten items or control items. Item-specific representations for remembered items were greater than both the forgotten items and control items, while the item-specific representations for forgotten items were not significantly different from the control items. *: *p* < 0.05; ***: *p* < 0.001; n.s.: not significant.

To further confirm that item-specific representations in the late time window were memory-related, we compared remembered items with non-memory control cues in an item type (remember vs. control) by item-specificity (WI vs. BI) two-way repeated measures ANOVA. The results also showed a significant interaction in a highly overlapped late time window (*p*_corr_ = 0.044, see Fig. S5). Post-hoc analysis [corrected for family-wise error rate (FWER)] on the late item-specific memory reactivation time window revealed that WI similarity was significantly greater than BI similarity for remembered items (*t*(29) = 3.01, *p*_FWER_ = 0.016), while there was no difference between WI and BI similarity for forgotten items (*t*(29) = -1.28, *p*_FWER_ = 0.637) or for control cues (*t*(29) = -0.95, *p*_FWER_ = 1.000) (Fig. 4B). Furthermore, item-specific representation (WI - BI similarity) for remembered items was greater than control items (*F*(1,29) = 6.31, *p* = 0.018), while there was no difference between item-specific representations for forgotten and control items (*F*(1,29) = 0.20, *p* = 0.657). Notably, results remained significant after controlling for trial pair number differences between WI and BI conditions and between memory and control items (Fig. S6). In addition, we found that pre-sleep testing did not modulate the item-specific representations (*p*_corr_ > 0.383, see Fig. S7).

Taken together, cue-elicited EEG activity in the early time window contained item-specific representations for both memory and control items, which may primarily reflect the effective processing of auditory stimuli during SWS. In contrast, the late time window item-specific representations emerged specifically for the post-sleep remembered items, which reflected the memory reactivation processes that support memory consolidation.

### Spindle activity coordinated item-specific memory reactivation for successfully remembered untested items

Previous studies suggest that TMR-based memory consolidation is tightly linked with post-cue spindle activity (*19, 27*). However, it remains unclear how spindle activity is related to memory reactivation and whether pre-sleep testing could affect the role of spindles in supporting memory reactivation. To address this question, we first detected discrete spindles on individual trials (Fig. 5A-B, see Materials and Methods) and then calculated spindle probability across all trials for each time point. We conducted pre-sleep testing (tested vs. untested) by subsequent memory (remember vs. forget) repeated measures ANOVA across individual time points and identified two significant clusters showing significant interaction effects (1^st^ cluster: 2344-3166 ms, *p*_corr_ = 0.002; 2^nd^ cluster: 3526-4118 ms, *p*_corr_ = 0.026; Fig. 5C). Post-hoc comparisons were performed by contrasting spindle probabilities across the two clusters. The results showed that, among tested items, the spindle probability for remembered items was not different from that for forgotten items (*t*(29) = -2.50, *p*_FWER_ = 0.147, Fig. 5D). In contrast, among the untested items, spindle probability for remembered items was greater than for forgotten items (*t*(29) = 4.50, *p*_FWER_ < 0.001). Moreover, post-sleep remembered untested items elicited higher spindle probability than both post-sleep remembered tested items (*t*(29) = 4.51, *p*_FWER_ < 0.001) and control items (*t*(29) = 5.69, *p*_FWER_ < 0.001, Fig. 5D), while no significant difference was found between the post-sleep remembered tested items and control items (*t*(29) = 1.42, *p*_FWER_ = 1.000). To ensure that the observed spindle activity difference in the late cluster was not due to the leakage of longer duration or delayed onset of the early spindles, we examined the duration and onset of spindles from the early (0-2 s) time window. The results revealed no significant interaction effect between memory and pre-sleep testing for either the duration or the onset of early spindles (all *p*s > 0.370, see Fig. S8). In addition, spindles that occurred in an extended time window of these two clusters (i.e., 2200-4200 ms) were significantly coupled to the up-state of SOs for all tested and untested items (all *ps* < 0.045) (see Fig. S9).

**Fig 5.**
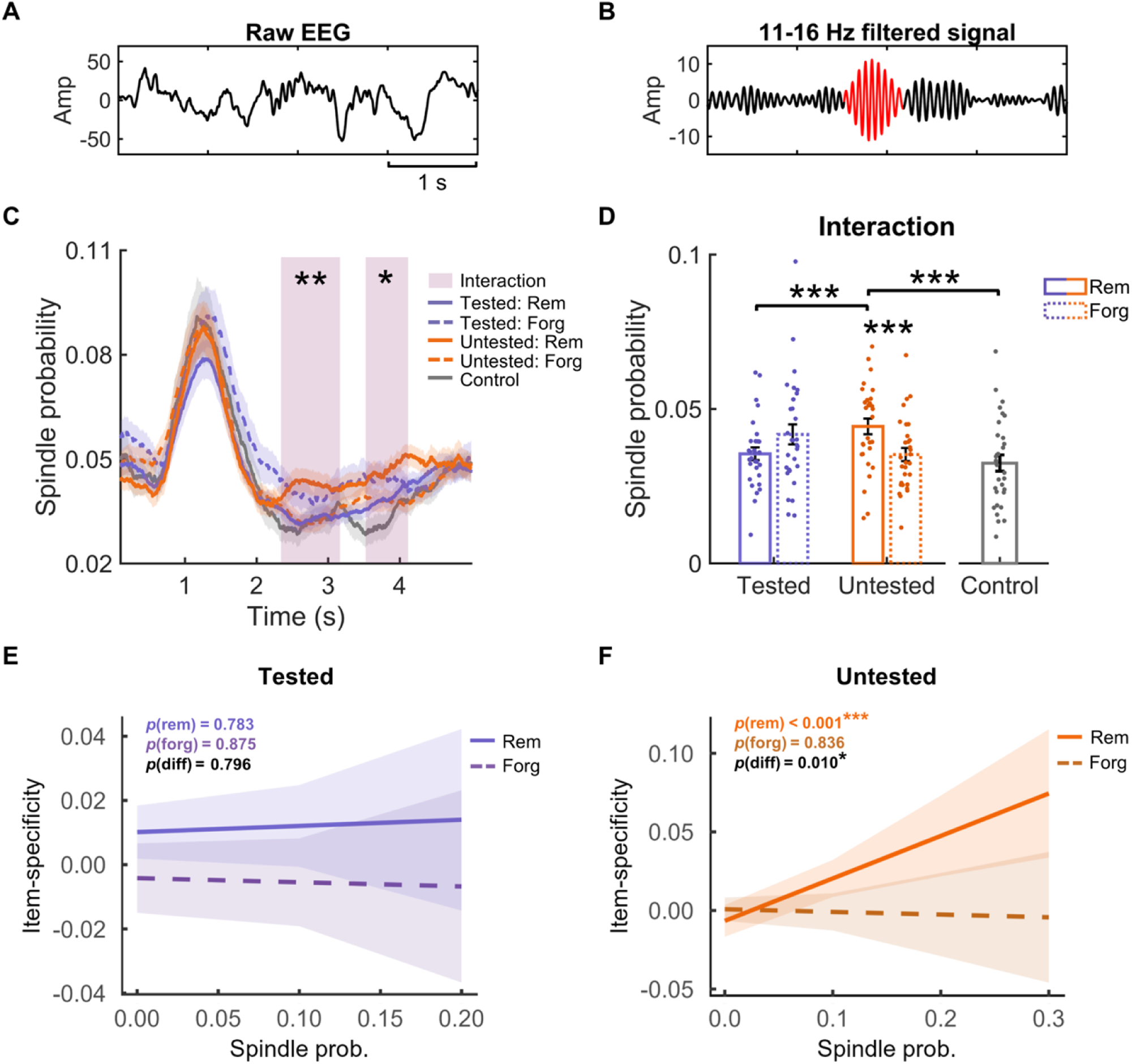
The relationship between spindles and item-specific memory reactivation in the overlapped late time window (i.e., 2500-2960 ms relative to cue onset). **(A)** An example of raw EEG data used for spindle detection. **(B)** EEG signals in (A) filtered between 11-16 Hz, with a spindle event highlighted in red. **(C)** Significant interaction effects between pre-sleep testing (tested vs. untested) and subsequent memory effect (remembered vs. forgotten items) on spindle probability in two clusters (2344-3166 ms and 3526-4118 ms, highlighted in shaded rectangles) following TMR cues. **(D)** Post-hoc analyses were performed on the averaged data across the two interaction clusters (shaded rectangle in C). For untested items, post-sleep remembered items elicited greater spindle probability than forgotten items. Moreover, the spindle probability for post-sleep remembered untested items was greater than that for both the post-sleep remembered tested items and control items. **(E)** Among tested items, the spindle probability did not predict the item-specific memory reactivation for either post-sleep remembered or forgotten items. **(F)** Among untested items, the spindle probability significantly predicted the item-specific memory reactivation for post-sleep remembered items but not for forgotten items. The effect for post-sleep remembered items was greater than that for forgotten items. *: *p* < 0.05; **: *p* < 0.01; ***: *p* < 0.001.

We next sought to establish the relationship between spindle activity and item-specific memory reactivation. Notably, increased spindles (2344-3166 ms and 3526-4118 ms in Fig. 5C) for post-sleep remembered untested items tended to co-occur in the late item-specific memory reactivation window (2500-2960 ms in Fig. 4A). Importantly, repeating the RSA using the spindle-related sigma power did not find any item-specific representations (*p*_corr_ > 0.111, Fig. S10), ruling out the possibility that late item-specific representations and spindles were different measures of the same phenomena. We then examined how spindles related to item-specific memory reactivation in the overlapping time window (i.e., 2500-2960 ms) for tested and untested items, respectively. Our linear mixed-effect model revealed that there were no significant effects for subsequently remembered or forgotten items among tested items (remember: *β* = 0.02, *t*(432) = 0.28, *p* = 0.783; forget: *β* = -0.01, *t*(258) = -0.16, *p* = 0.875, Fig. 5E). However, among untested items, cue-elicited spindle probability significantly and positively predicted item-specific memory reactivation for subsequently remembered items (*β* = 0.26, *t*(239) = 3.678, *p* < 0.001, Fig. 5F) but not for forgotten items (*β* = -0.02, *t*(472) = -0.21, *p* = 0.836). Moreover, among untested items, the effect for remembered items was significantly greater than that for forgotten items (*β* = 0.29, *t*(716) = 2.57, *p* = 0.010). These results suggest that sleep spindles play a critical role in supporting item-specific memory reactivation and contribute to memory consolidation, particularly for untested items.

## Discussion

We asked fundamental yet unanswered questions in exogenous memory reactivation: how individual sensory memory cues reactivate the corresponding memory traces during slow-wave sleep and how such item-specific reactivation led to consolidation. Using targeted memory reactivation (TMR) and multivariate representational similarity analyses (RSA), we found highly specific item-level representations, which went beyond prior studies showing neural reinstatement effects of broader, categorical types (*19–21*). Specifically, both memory cues and non-memory control cues first elicited item-specific representations, likely reflecting the successful sensory processing of individual auditory cues. Item-specific memory reactivation emerged in a late time window that was only present for memory cues, which contributed to post-sleep memory retention. Critically, increased spindles during this late memory-specific window preferentially supported item-specific memory reactivation only among pre-sleep untested items, highlighting that testing can be an important factor that modulates sleep-based memory consolidation (see Fig. 6).

**Fig 6.**
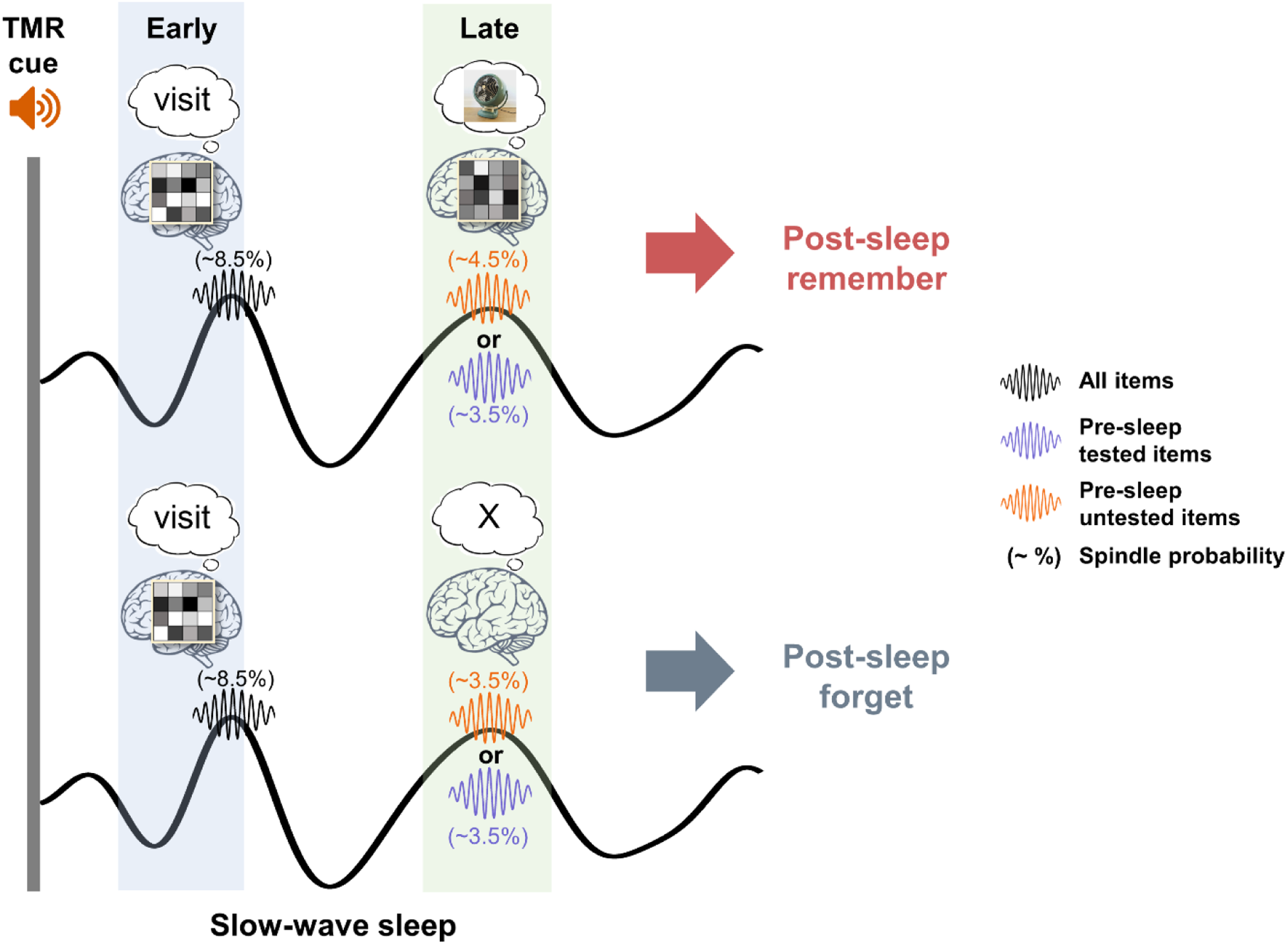
Schematic depiction of item-specific memory reactivation following TMR cues promotes memory consolidation. Sensory memory cues trigger early item-specific representations during SWS, predominantly reflecting sensory processing. In contrast, item-specific memory reactivation and spindle activity within slow-oscillation up-states in a later time window contribute to memory retention for pre-sleep untested items.

Even during SWS, the brain preserves its capacity to process external sensory information, which is a prerequisite for sensory memory cues to reactivate its associated memories (*23*). Cross-species studies have consistently demonstrated that the primary auditory cortex neuronal responses are well preserved during sleep, with little modulation by different vigilance or consciousness states (*24, 44, 45*). Humans research further shows that the sleeping brain can process meaningful auditory stimuli (*46–49*), which are manifested by increased EEG power in both the low-frequency and sigma band immediately following auditory stimuli (*41, 42*). Consistent with these previous studies, we found that both the memory cues and non-memory control cues enhanced low-frequency activities (1-10 Hz), followed by the extended sigma band activities (11-18 Hz) within approximately the first 2 s post-cue onset. Beyond EEG spectral power changes, our results further revealed that the cue-elicited EEG contained highly specific, item-specific representations in this early cluster. Moreover, this early item-specific representation was identified for all memory cues regardless of post-sleep memory performance and for control cues that were not paired with any prior learning. These early, item-specific representations may thus reflect the initial processing of individual auditory cues.

While previous studies often found greater EEG power in response to memory cues than control cues (*19, 50*), we did not find significant differences. This discrepancy might be caused by the novelty of the control cues: whereas in these previous studies, control cues were not presented before the TMR session; here, we presented all memory and control cues at least twice during the pre-learning familiarization phase. One possibility is that such familiarization would evoke strong K-complex activities in the sleeping brain (*51–53*), thus minimizing the differences between memory and control cues. Notably, these early EEG responses elicited by the cues predicted the subsequent behavioral TMR effects, despite almost unitary presentations of auditory cues across participants. These findings are consistent with previous research that has linked this early cue-elicited EEG power with behavioral TMR effects (*29, 50, 54, 55*). Moreover, our results showed that even the increased EEG power elicited by non-memory control cues could predict the early item-specific representations of the individual memory cues. Our findings provided new insights that how early EEG responses reflect sensory processing of memory cues, which paves the pathways for successful memory reactivation in later time windows.

Indeed, we found compelling evidence that item-specific representations in a late time window reflected the item-specific memory reactivation. Specifically, post-sleep remembered items exhibited greater item-specific representations than forgotten items and control cues in this late cluster. The differences in item-specific representations cannot be exclusively attributed to different levels of familiarity with sensory cues, given that the cues for post-sleep remembered and forgotten items were repeated the same number of times during pre-sleep learning. In addition, the direct comparisons between the item-specific representations of forgotten items and control items did not yield significant differences. Notably, this late item-specific memory reactivation window overlaps with the spindle refractory period, highlighting that this period is instrumental for memory reprocessing during NREM sleep (*25*).

Item-specific memory reactivation following TMR cues could be achieved via a cortical-hippocampal-cortical loop (*56–58*): effective early processing of auditory cues in the neocortex reactivate their associated memory traces that are temporarily stored in the hippocampus (*11, 59–61*). The reactivated hippocampal memory traces then trigger the reinstatement of the neocortex-dependent memory representations, which may follow the same principle of pattern completion as during wakefulness retrieval (*62, 63*). This cortical-hippocampal-cortical loop during NREM sleep relies on the timely coupling of thalamocortical spindles to the up-state of slow oscillations in both humans and rodents, serving as the foundation for systems memory consolidation (*3, 64–67*). Critically, our results showed that item-specific memory reactivation, supported by spindles, during the late window predominantly contributes to memory consolidation. Moreover, the accompanying spindles were preferentially coupled to the up-state of slow oscillations, which may have facilitated cross-regions interactions for reactivated memories, particularly the pre-sleep untested items, to be consolidated (*68–70*). A more precise delineation of the spindle-SO-coordinated cross-regional interaction awaits future investigations using methods affording both high spatial and temporal resolutions (e.g., intracranial EEG recordings).

Despite spindles’ importance in supporting item-specific memory reactivation, pre-sleep testing modulated this spindle-mediated consolidation process. Given that pre-sleep testing largely boosted memory performance relative to untested items in the current study, it is possible that pre-sleep tested items already underwent retrieval-induced fast consolidation before sleep (*30*). Thus, spindles may preferentially support item-specific memory reactivation for untested memories. Corroborating this possibility, previous studies have shown that strong memories did not further benefit from subsequent spontaneous sleep (*71*) or sleep TMR (*34*). Moreover, sleep spindles would preferentially consolidate weak over strong memories (*36*), and TMR promoted consolidation for memories with low (*33*) or moderate (*34*) pre-sleep accuracy. Consistent with this idea, our results also showed cue-elicited early EEG power was associated with TMR effects only for pre-sleep untested items.

Our study did not find a significant TMR behavioral effect. A few possibilities might account for the absence of TMR effects. First, as mentioned above, TMR may be less effective for strong memories (*33, 34*). Here, participants may overlearn the word-picture pairs via repeated encoding and mental rehearsal during the pre-sleep learning, which may contribute to fast consolidation and thus weaken the TMR effects (*30*). In addition, the pre-sleep recognition task might serve as the feedback after the pre-sleep test, which further strengthens the pre-sleep memory for tested items and attenuates the TMR effect. Second, given that TMR effects are highly sensitive to memory measures (*10*), more precise measurements are desirable to detect the TMR effects, such as the error distance measures used in previous TMR research (*11, 12*). Future word-picture or naturalistic episodic memory paradigms could also include verbal recall to test how TMR impacts perceptual and conceptual details.

In conclusion, the identification of item-specific memory reactivation and the segregated processing windows contributed to the mechanistic understanding of how exogenous memory reactivation unfolds during NREM sleep. Effective processing of individual TMR cues would drive the reactivation of item-specific memory representations for consolidation, supported by spindles in a later time window. These findings may ignite new development of sleep-based memory editing techniques (*72*) in perturbing individual memories: by targeting the underlying neural activity at the critical memory reactivation time window, techniques can either strengthen newly acquired knowledge or selectively weaken maladaptive memories.

## Materials and Methods

### Participants

Thirty healthy, right-handed participants were included in the analyses (23 females, mean ± SD age, 22.37 ± 2.94 years). To obtain reliable results in the EEG power spectral analysis as well as the multivariate representational similarity analysis for both the memory items and control items, we set a minimum requirement of 5 rounds of TMR cueing (i.e., 20 trials). Thus, seven additional participants failed to meet this criterion due to insufficient or unstable N2 and N3 sleep were excluded from the analysis. All included participants had normal or corrected-to-normal vision. Participants were pre-screened on sleep by using the questionnaires including the Pittsburgh Sleep Quality Index (PQSI) and the Insomnia Severity Index (ISI). They all reported overall good sleep quality and had not taken any medicines to aid sleep in the past month before the experiment. All participants did not suffer from any neurological or psychiatric disorders. The study was approved by the Research Ethics Committee of the University of Hong Kong. All participants gave written informed consent prior to the experiment.

### Stimuli

A total of 96 two-character Chinese verbs and 96 target pictures were used in the experiment. Each verb was randomly paired with a complex visual picture, resulting in 96 cue (word)-target (picture) pairs. The central element of each picture was from one of four categories (i.e., animals, electronic devices, plants, and transportation tools), with 24 pictures in each category. For each picture, a highly similar picture was also selected and served as a lure in the old/new recognition task. For each unique picture, the target and lure were randomly assigned among participants. For verbs, visually presented verbs were only used in the familiarization phase, while aurally presented verbs were used throughout the entire experiment. Four additional two-character Chinese verbs were presented during the familiarization phase but were never paired with any pictures. These verbs served as control cues in the TMR. Auditory sounds of the verbs were generated using the Text-To-Speech of iFLYTEK, with an average duration of 631.70 ms (SD: 55.40 ms).

### Procedure

All participants arrived at the sleep lab around 8:30 pm. Participants completed the following tasks in order: a vigilance task, a stimuli familiarization task, a cue-target associative learning task, and a pre-sleep memory test. Participants then proceeded to sleep (12 am to 8 am the next morning), wherein TMR was administered during slow-wave sleep. After ∼30 min upon wakening, participants’ vigilance levels were assessed again, followed by the post-sleep memory test. All behavioral tests were conducted via using the psychopy (version 2020.2.10).

#### Psychomotor vigilance task

Participants’ vigilance levels were assessed using the Psychomotor Vigilance Task (PVT), right after participants arrived at the sleep lab and the next day morning. During the vigilance task, a fixation was presented on the center of the screen with a jitter duration in the range of 2-10 s. Then, the fixation was replaced by a counter counting incrementally from 0 in 10 ms increments. Participants were instructed to stop the counter by pressing the space bar immediately after they detected the change of fixation to the number. Response time that appeared on the screen would serve as feedback on the performance. This task lasted for 5 minutes.

#### Cue word and picture familiarization task

The familiarization task consists of two sessions, a cue word-familiarization session, and a picture-familiarization session. In the cue word-familiarization session, each trial started with a 0.3 s fixation, followed by a 0.5 s blank screen. Afterward, a verb was visually presented on the center of the screen for 2 s, accompanied by its verbalization from the speaker. Participants judged whether the spoken verbs were clear and recognizable by pressing a button. All 100 verbs (96 verbs in the word-picture pairs and 4 verbs as control cues) were randomly presented during this stage, with each verb being presented twice. In the picture familiarization session, each trial started with 0.3 s fixation, followed by a 0.5 s blank screen. Afterward, a picture and its label (e.g., for a panda picture, the label would be “panda”) were presented on the screen for 2 s. Participants indicated whether they were familiar with the picture and its name by pressing a button. Each of the 192 pictures (96 targets + 96 lures) was presented twice. For items that participants indicate unfamiliar will be presented for another two rounds. At the end of the task, participants indicated that all stimuli were clearly recognizable and familiar.

#### Word-picture associative learning task

Participants learned 96 word-picture pairs. Each learning trial consisted of three phases, an encoding phase, a maintenance phase, and a vividness rating phase. During the encoding phase, following a 0.3 s fixation and a black screen jittering between 0.9-1.5 s, participants viewed a picture presented in the center of the screen for 2 s while hearing the spoken verb from the speaker. Participants were instructed to pay attention to the details of the picture while memorizing the verb-picture associations during the encoding phase. During the maintenance phase, the picture disappeared, and participants were asked to maintain the picture in their minds as vividly as possible for 3s while hearing the spoken verb again. In the vividness rating phase, participants indicated the vividness of the mental image of the picture on a 1 (not vivid at all) -4 (very vivid) point Likert scale by pressing one of four buttons. The learning task consisted of three blocks, with each block containing 32 unique verb-picture pairs that were repeated three times. To reduce the recency effect, participants completed a ∼5 min math task after the learning task.

#### Pre-sleep memory test

To understand whether the pre-sleep testing alters TMR effects on memory consolidation, participants were tested on half of the pairs during the pre-sleep test (i.e., 48 pairs, with 12 pairs from one of the four picture categories). This test consisted of a cued recall task and a cued recognition task.

In the cued recall task, each trial started with a 0.3 s fixation, followed by a blank screen (0.9-1.5 s). The spoken verb was played, prompting participants to report whether they could “remember” or “forget” the corresponding picture. This stage was self-paced so that participants had enough time to recall. Immediately following this “remember” or “forget” response, participants were further asked to report the category of the picture by pressing one of four buttons, with each button indicating one of four categories within 2 s.

To further encourage participants to remember the detailed picture content, instead of the category information, we further ask participant to perform a cued-recognition task. In this recognition task, the same half of the pairs were tested. Specifically, each trial started with a fixation (0.3 s) and was followed by a blank screen (0.9-1.5 s). Participants next saw the picture presented on the center of the screen while hearing the spoken verb. Participants were asked to indicate if the picture was the same picture paired with the verb during the previous learning task by pressing the “Yes” or “No” button. Among 192 recognition trials, 48 trials showed identical verb-picture associations from the learning task, and 48 trials showed verb-lure picture associations. The remaining 96 trials consisted of 48 mixed verb-picture associations and 48 mixed verb-lure associations with cue verbs and pictures (or lures) corresponding to different learning trials.

#### TMR during NREM sleep

To counterbalance the pre-sleep memory performance between cued and uncued items, we selected half of the remembered and half of the forgotten pairs (based on the category report performance in the cued recall task) of the tested items as cued items during TMR. In addition, we randomly selected half of the untested items to be cued in the TMR. Picture categories were balanced across cued versus non-cued conditions. Thus, the TMR session contained 48 verbs as cues, with additional 4 verbs that were not paired with any pictures as control cues. Half of the cues were from tested trials, with the remaining half from untested trials. The control cues served as the baseline to examine retrieval-specific neural activity. Therefore, there were 52 unique sound cues played during sleep.

During the nocturnal sleep, white noise was played in the background throughout the night, with an intensity of ∼ 45 dB, measured by a sound-level meter placed at the same position where participants laid their heads on the pillow. Experienced experimenters monitored the EEG signals and visually identified signature EEG events characterizing different sleep stages (e.g., spindles, K-complex, slow oscillations). Upon detecting stable slow-wave sleep, the experimenter would begin the TMR, which occurred ∼50.04 min (SD: 37.21min) after the start of the sleep phase. On average, participants were presented with 8.92 (SD: 3.24) rounds of TMR. In each round of TMR, 52 verb cues were randomly presented with an inter-stimuli interval of 5 ± 0.2 s. After each round of cueing, the order of the TMR cues was shuffled and replayed again. Each round was separated by 30 s. The TMR session continued as long as the participants were in slow-wave sleep in the first 3-4 hours of the nocturnal sleep. Cueing was stopped immediately when participants showed signs of micro-arousals, awakening or changed to N1 sleep or rapid eye movement (REM) sleep. Cueing was resumed after the participants returned to stable slow-wave sleep.

#### Post-sleep memory test

Approximately 30 minutes after awakening and after the PVT test, participants were tested on all 96 pairs. Similar to pre-sleep memory tests, this test included the cued recall and cued recognition tasks. In addition, there was a closed-eye mental retrieval task between these two tasks, in which participants were asked to keep their eyes closed and relax while each of the verbs was randomly played via the speaker (ISI = 5 ± 0.2 s). The current study focuses on TMR-based memory consolidation process. Therefore, the post-sleep closed-eye mental retrieval data will report elsewhere.

### EEG recording and preprocessing

We collected the EEG data throughout the experiment except for during the familiarization and PVT tasks. EEG data were recorded using the amplifier from the eego system (ANT neuro, Netherlands, https://www.ant-neuro.com) with a sampling rate of 500 Hz from 61 channels (waveguard EEG caps) that were mounted in the International 10-20 system. Additionally, there were two electrodes placed on the left and right mastoids, respectively, and one electrode was placed above the left eye for the EOG measurements. Online EEG recordings were referenced to the default reference channel (i.e., CPz). For sleep monitoring, another two electrodes were placed on both sides of the chin to measure the EMG using the bipolar reference. EEG data collection was started after the impedance of all electrodes was lower than 20 KΩ.

Sleep EEG data preprocessing was performed using EEGLAB (https://sccn.ucsd.edu/eeglab/) and Fieldtrip toolboxes(*73*), as well as in-house code that was implemented in MATLAB (MathWorks Inc.). EEG data were first notch filtered at 50 ± 2 Hz, and then bandpass filtered between 0.5 and 40 Hz. Then continuous sleep EEG data were segmented into 15 s epochs, i.e., form -5 s to 10 s relative to the onset of TMR cues. This long epoch was used to eliminate the edge effect in the later time-frequency analysis. Our main interesting time windows for the sleep data are from 0 to 5 s relative to the TMR cue onset. Bad epochs are marked based on visual inspection and rejected from further analysis. Bad channels were marked and interpolated using spherical interpolation in EEGLAB. Afterward, EEG data were re-referenced to the average of the artifact-free data.

### Time-frequency analysis

We performed time-frequency transformation on the preprocessed EEG data, using the complex Morlet wavelets (six cycles). Spectral power was extracted from the frequency range of 1-40 Hz, with a step of 1 Hz and with the time of interest range of [-1 5 s] relative to the TMR cue onset. The power was down-sampled to 100 Hz after the time-frequency transformation. We normalized the power data within each frequency bin and each channel by first subtracting the mean power in the baseline time windows ([-1 -0.5s] relative to the TMR cue onset) and then dividing the same baseline mean power. Finally, we re-segmented the power data into the 5s epochs (i.e., [0 5s] relative to the TMR cue onset).

### Spindle detection

Individual spindles were detected during the TMR periods following previous studies(*27*). Specifically, artifact-free EEG data during the TMR periods were first bandpass filtered between 11-16 Hz by using the 4^th^ order two-pass Butterworth filter. Next, the root-mean-square (RMS) values were calculated for each time point with a moving window of 400 ms. Third, the spindle amplitude criterion is defined as the mean + 1.5 SD of the RMS signal. Sleep spindles were detected if the RMS signal consecutively exceeded the amplitude criterion for a duration of 0.5 to 3 s. Previous studies indicated that spindle activity was prominent over anterior-posterior midlines(*74*). To this end, spindle detection was performed for all seven midline EEG channels (i.e., “FPz”, “Fz”, “FCz”, “Cz”, “Pz”, “POz”, and “Oz”), separately. We then assign the spindle value of 1 to the time points where a spindle was detected and 0 otherwise. The spindle probability was computed as the mean spindle values across trials for each time point in the time range of [0 5 s] relative to the TMR cue onset and then averaged across all midline channels.

### Slow-oscillation detection

The detection of slow oscillations (SOs) was performed on the artifact-free EEG data during the TMR periods. We first bandpass filtered the EEG data between 0.3 and 1.25 Hz using the two-pass FIR filter, with the order equalling the three cycles of the low-frequency cut-off. We then detected the zero-crossings in the filtered signal, and the event durations were calculated as the temporal distance between two successive positive-to-negative zero-crossings. SO was detected if the peak-to-peak amplitude was greater than the 75% percentile of the absolute amplitude of the filtered signal(*75*) and the event duration was between 0.8 s and 2 s. SOs detection was performed on the Fz, where the amplitude of SOs was prominent according to previous studies(*76*).

### SO-spindle coupling

For TMR trials that showed both the spindle and SOs in the time windows (i.e., 2500-2960 ms), which showed significant interactions of item-specificity and post-sleep memory effect, we then performed the SO-spindle coupling. We first filtered the SO-spindle trials in the SO frequency range (i.e., 0.3-1.25 Hz). We then applied the Hilbert transform to the filtered data to extract the instantaneous phase of the SOs. To obtain the amplitude of the spindle activities, we filtered the SO-spindle trials in the spindle frequency range (i.e., 11-16 Hz) and then applied the Hilbert transform to the filtered data to obtain the instantaneous amplitude of the spindle activities on an extended time window (i.e., [2.2-4.2 s] relatively to TMR cue onset to ensure at least half of the SOs cycle was included). We detected the preferred SO phase, which is concurrent with the maximal spindle amplitude across all trials in each participant, and then tested the distribution of the preferred SO phases across participants against the uniform distribution using the Rayleigh test (CircStat toolbox(*77*)).

### Representational similarity analysis

To extract item-specific representations, we performed representational similarity analysis (RSA) between every two clean TMR trials that originated from different TMR rounds. Specifically, the within-item similarity was calculated as the whole-brain EEG cortical pattern similarity between the same TMR cues; and the between-item similarity was calculated as the pattern similarity between different TMR cues. Thus, contrasting within-item similarity versus between-items similarity enables us to examine whether there are item-specific representations following the TMR cues.

To characterize the item-specific representations across time, RSA was performed on the preprocessed raw EEG using sliding time windows of 500 ms, with an incremental step of 10 ms, resulting in 250*61 (time points * channels) features in each time window. Then, for each time window, we calculate the similarity between vectorized features of every two trials that were from different TMR rounds using Spearman’s corrections. All the correlation values were Fisher *Z*-transformed before further statistical analysis.

## Statistics

For the behavioral data, we conducted a two-way ANOVA with TMR effects (cued vs. uncued) and pre-sleep testing (tested vs. untested) as repeated measures in analyzing the post-sleep memory performance. We also performed a two-way ANOVA with TMR and time (pre-sleep vs. post-sleep) as repeated measures in analyzing the memory performance for tested items. The paired sample *t*-tests were employed to examine the differences between two specific experimental conditions (e.g., memory performance between pre-sleep tested items and post-sleep tested items). The relationship between EEG activities during the TMR period and post-sleep memory performance is assessed by the robust linear regression model, which is less likely to be affected by potential outliers(*78*).

For the EEG data, multiple comparisons across consecutive time windows were corrected using the cluster-based nonparametric statistical tests in MATLAB(*79*). Specifically, statistical tests (e.g., *t*-test) were performed between conditions (e.g., WI vs. BI or tested vs. untested) in individual time (or time-frequency) windows. Adjacent time (or time-frequency) windows with statistical values exceeding a threshold (*p* < 0.05) were combined into contiguous clusters. Cluster-level statistics were computed using the sum of the *t* values within a cluster. To test the significance of the time (or time-frequency) cluster, a distribution of cluster-level statistics under the null hypothesis was constructed by randomly permuting condition labels 1000 times, and the maximum cluster-level statistic in each permutation was extracted. If no significant cluster was found for a permutation, a value of 0 was assigned for that permutation. The nonparametric statistical significance of a cluster was then obtained by calculating the proportion of cluster-level statistics in the distribution under the null hypothesis that exceeded the empirical cluster-level statistics. For an identified cluster, we also performed a two-way ANOVA with item-specificity (WI vs. BI similarity) and pre-sleep testing as repeated measures and a two-way ANOVA with item-specificity and subsequent memory as repeated measures. The statistical significance level for all the analyses is set as *p*-value < .05 or corrected *p*-value (*p*_corr_) < 0.05.

## Acknowledgments

**General:** We thank Lingqi Zhang and Zexuan Mu for their help with the data collection.

## Author contributions

J. L., and X. H. conceived the study and designed the experiment. J. L., and M. Z. conducted the experiment. J. L., T. X., D. C., Z. Y., J. W. A., T. L., and X. H. contributed to data analyses. J.L., X. H. and T.L. drafted the manuscript, with critical feedback from J.W.A. All authors contributed to the revision of the manuscript.

## Data and materials availability

All data needed to evaluate the conclusions in the paper are present in the paper and/or the Supplementary Materials. All data that support the findings of this study are available in the Open Science Framework (https://osf.io/knb42/?view_only=04d1d0d6deb24c69b34cacce73ed6f03).

