## Supplementary Materials for "Item-specific memory reactivation during sleep supports memory consolidation in humans"

Jing Liu *et al.*

**This PDF file includes:**

Figs. S1 to S10

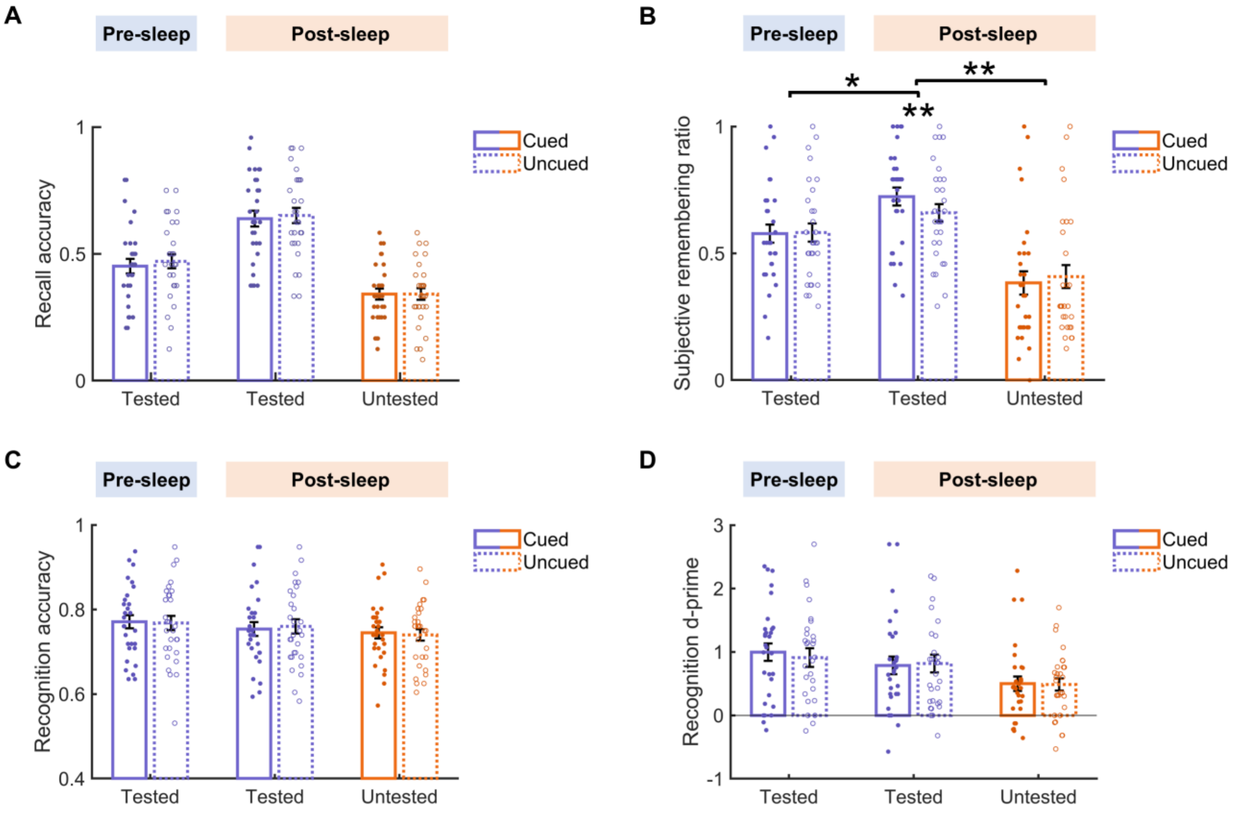


**Fig S1. Behavioral results. (A)** Category-report accuracy during pre- and post-sleep cued-recall tasks. In the cued recall task, participants were presented with auditory cue words and were asked to report their subjective recollection of associated pictures along with their respective categories. Category-report accuracy was higher in the post-sleep test compared to the pre-sleep test for tested items, irrespective of TMR cueing ($t(29) = 9.85$, $p_{FWER} < 0.001$). Moreover, memory accuracy for tested items was significantly higher than that for untested items in the post-sleep test ($t(29) = 13.67$, $p_{FWER} < 0.001$). A repeated measures ANOVA with TMR (cued vs. uncued) and pre-sleep testing (tested vs. untested) as factors on post-sleep category-report accuracy revealed neither a significant interaction effect ($F(1,29) = 0.15$, $p = 0.700$) nor a significant main effect of TMR (cued vs. uncued, $F(1,29) = 0.11$, $p = 0.739$). In addition, for pre-sleep tested items, there was no significant TMR (cued vs. uncued) by time (pre-sleep vs. post-sleep) interaction effect ($F(1,29) = 0.11$, $p = 0.745$). **(B)** Subjective remembering during pre- and post-sleep cued-recall tasks. Subjective remembering was quantified by the ratio of trials that participants reported "remembering" during the cued-recall test regardless of the following category report accuracy. The results found that subjective remembering for the tested items in the post-sleep test was greater than that in the pre-sleep test, as well as greater than the untested items in the post-sleep test, irrespective of TMR cueing (Both $ps_{FWER} < 0.001$). Further analysis revealed a significant TMR by pre-sleep testing (tested vs. untested) interaction effect in the post-sleep test and a significant TMR (cued vs. uncued) by time (pre- vs. post-sleep) interaction effect for tested items ($ps < 0.022$). Simple-effects analysis revealed that both interaction effects were driven by greater subjective remembering for cued than uncued items among the tested items during the post-sleep test ($t(29) = 3.80$, $p_{FWER} = 0.002$). **(C)** Recognition accuracy. Recognition accuracy for tested items in the pre-sleep test was significantly greater than that for the untested items in the post-sleep test ($t(29) = 3.13$, $p_{FWER} = 0.012$). However, there was no significant difference in the accuracy between tested items in the pre-sleep test and the post-sleep test, nor between tested and untested items in the post-sleep test ($p_{FWER} > 0.192$). Further analyses revealed no significant TMR by pre-sleep testing interaction effect in the post-sleep test, nor a significant TMR by time interaction effect for tested items ($p$s > 0.351). **(D)** Recognition d-prime. The d-prime values for tested items in both the pre-sleep and post-sleep tests were significantly higher than untested items ($p$s$_{FWER} < 0.004$). However, there was no significant difference in d-prime values for tested items in the pre-sleep and post-sleep tests ($t(29) = 2.28$, $p_{FWER} = 0.091$). Further analyses revealed no significant TMR by pre-sleep testing interaction effect in the post-sleep test, nor a significant TMR by time interaction effect for tested items ($p$s > 0.314). *: $p < 0.05$; **: $p < 0.01$.

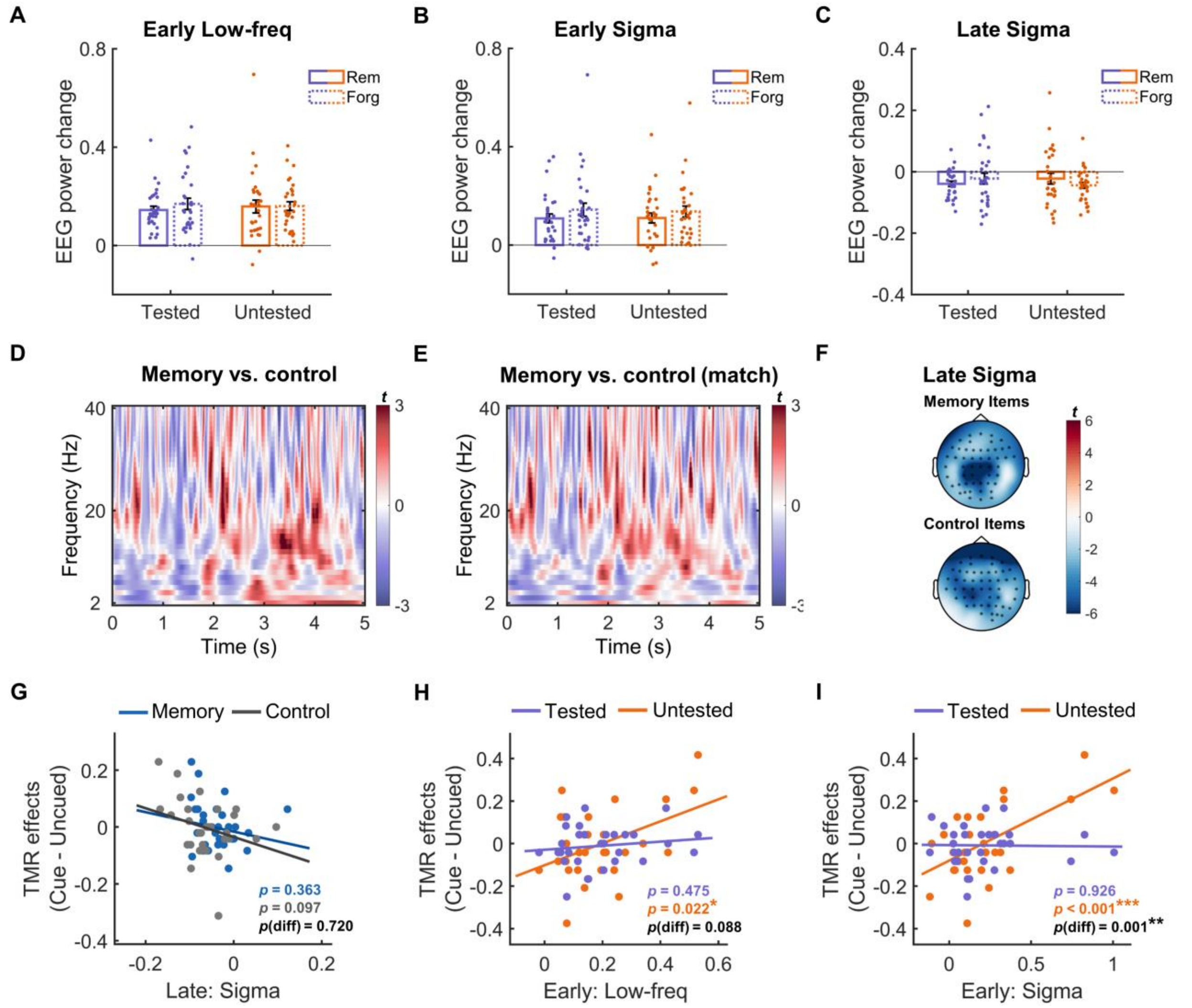


**Fig S2. Auditory cue-elicited EEG power and its association with post-sleep TMR effects.** **(A-C)** Pre-sleep testing (tested vs. untested) by memory (remember vs. forget) two-way repeated measures ANOVA on memory cue-elicited EEG power showed no significant interaction effects in either early clusters (i.e., low-frequency and sigma band clusters) or the late cluster (i.e., reduced sigma power cluster) (all *p*s > 0.102). **(D)** No significant difference in auditory cue-elicited EEG power between memory items and control items. ($p_{corr} > 0.294$). **(E)** After matching the trial number between memory cues and control cues, the difference in cue-elicited EEG power remained non-significant ($p_{corr} > 0.441$). **(F)** Topography plots for the cue-elicited reduced sigma band power in the late cluster for memory cues and control cues, respectively. **(G)** Neither memory cue nor control cue-elicited late sigma band power predicted post-sleep TMR effects (memory cue: adjusted $R^2 = -0.004$, $p = 0.363$; control cue: adjusted $R^2 = 0.06$, $p = 0.097$;). Control cue-elicited EEG power in both the low-frequency range **(H)** and sigma band **(I)** in the early cluster significantly positively predicted the post-sleep TMR effects for untested items (Low-frequency: adjusted $R^2 = 0.14$, $p = 0.021$; Sigma: adjusted $R^2 = 0.32$, $p < 0.001$), but not for tested items (Low-frequency: adjusted $R^2 = -0.02$, $p = 0.475$; Sigma: adjusted $R^2 = -0.03$, $p = 0.926$). *: $p < 0.05$; **: $p < 0.01$; ***: $p < 0.001$.

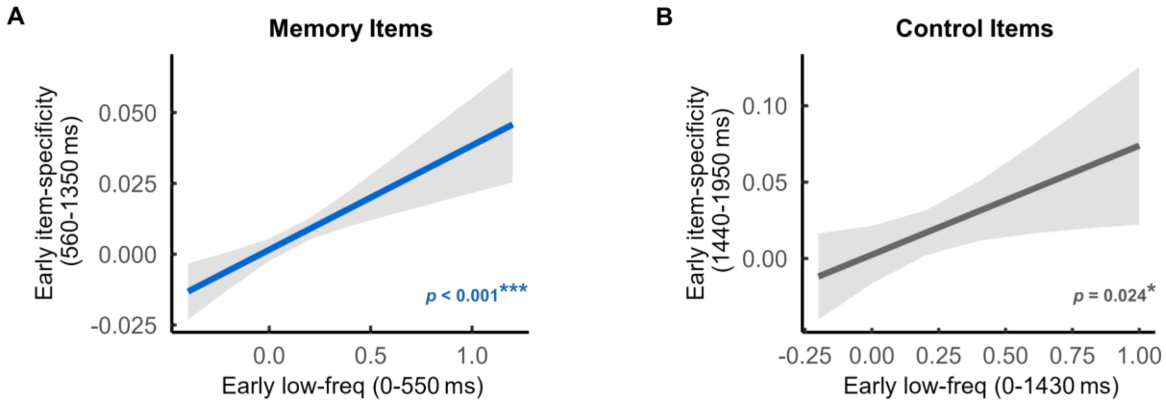


**Fig S3. Auditory cue-elicited low-frequency EEG power in the very early time window predicted item-specific representations in the subsequent non-overlapped early clusters.** To investigate the link between auditory cue-elicited EEG power and the identified item-specific representations in the early time window while controlling for the possible influence of EEG power on item-specific representations, we utilized EEG power from an earlier time window that did not overlap with the time windows demonstrating item-specific representations. **(A)** Memory cue-elicited low-frequency activities in the very early time window, i.e., 0-550 ms post-cue, significantly and positively predicted the memory cue-elicited item-specific representations in the 560-1350 ms cluster ($\beta = 0.04$, $t(1273) = 3.92$, $p < 0.001$). **(B)** Control cue-elicited low-frequency activities in the 0-1430 ms time window significantly and positively predicted the control cue-elicited item-specific representations in the 1440-1950 ms cluster ($\beta = 0.07$, $t(117) = 2.28$, $p = 0.024$). *: $p < 0.05$; ***: $p < 0.001$.

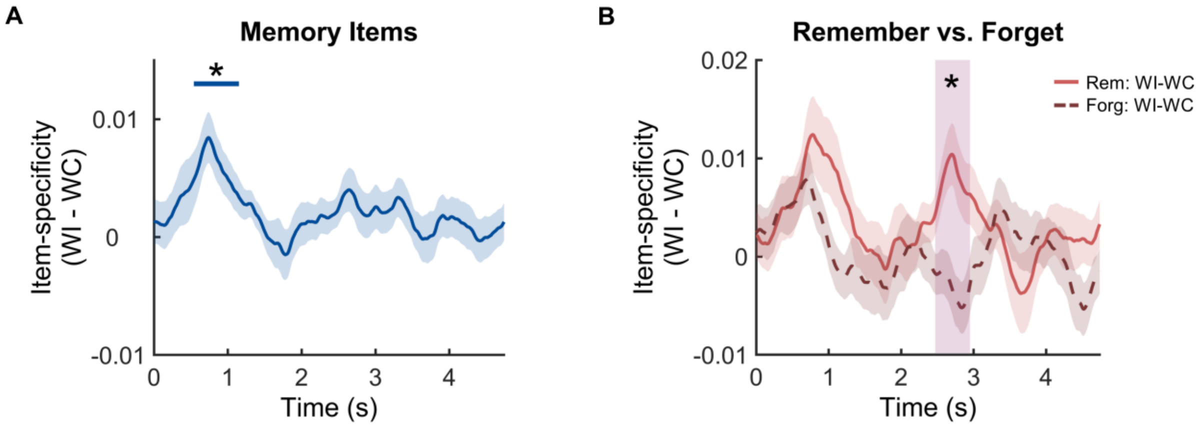


**Fig S4. Item-specific representations for memory items after constraining the between-item similarity to trial pairs with pictures from the same category (i.e., within-category similarity).** The within-item (WI) similarity was contrasted with the within-category (WC) similarity. **(A)** Item-specific representations were identified in a 550-1160 ms time window after cue onset across all memory items ($p_{corr} = 0.013$). **(B)** Greater item-specific representations for post-sleep remembered items than that for forgotten items in the 2480-2960 ms time window after cue onset ($p_{corr} = 0.027$). These results are highly consistent with Fig. 3C in the main text. *: $p < 0.05$.

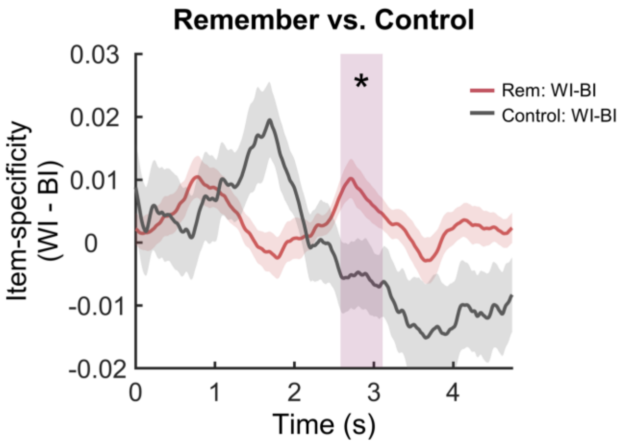


**Fig. S5. Greater item-specific representations for post-sleep remembered items than control items in the late time window.** An item type (remembered vs. control item) by item-specificity (WI vs. BI similarity) repeated measures ANOVA revealed a significant interaction effect cluster in a late time window (2590-3120 ms, $p_{corr} = 0.044$). Post-hoc analysis on this interaction cluster showed that WI similarity was significantly greater than BI similarity for remembered items ($p_{FWER} = 0.019$), while not for control items ($p_{FWER} = 0.513$). *: $p < 0.05$.

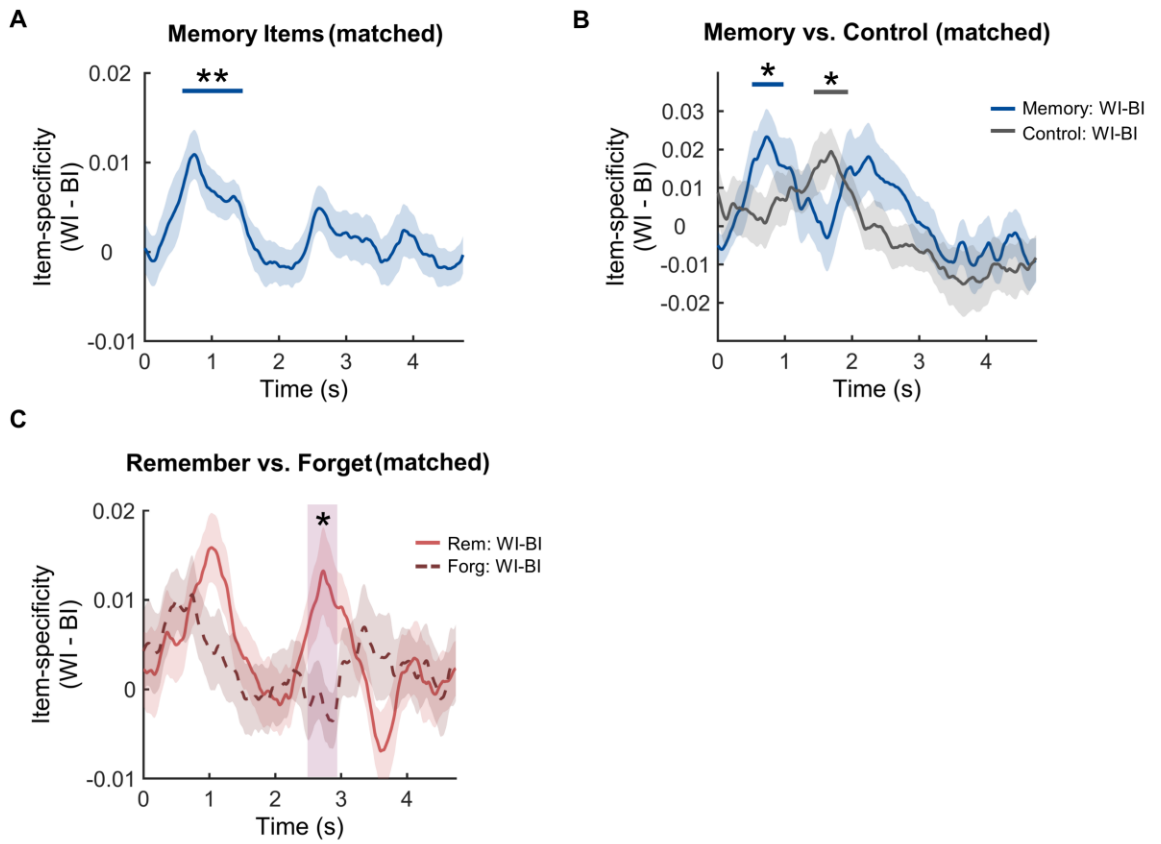


**Fig S6. Item-specific representations after controlling the trial pair number between conditions. (A)** After matching the trial pair number between WI similarity and BI similarity conditions, we still identified a significant cluster that showed item-specific representations for memory items (570-1470 ms, $p_{corr} = 0.003$). **(B)** After matching the trial pair number of the WI and BI similarity for memory items with that for the control items, we still found a significant cluster that showed item-specific representations for memory items (520-990 ms, $p_{corr} = 0.038$). An item type (memory vs. control items) by item-specificity (WI vs. BI) repeated measure ANOVA did not reveal any significant interaction effect cluster ($p_{corr} > 0.224$). **(C)** Greater item-specific representations for post-sleep remembered items than forgotten items in a late time window (2500-2950 ms, $p_{corr} = 0.035$) after matching the trial pair number between post-sleep remember and forget conditions. These results indicate that item-specific representations persisted after matching the trial pair number used in the representational similarity analysis. *: $p < 0.05$; **: $p < 0.01$.

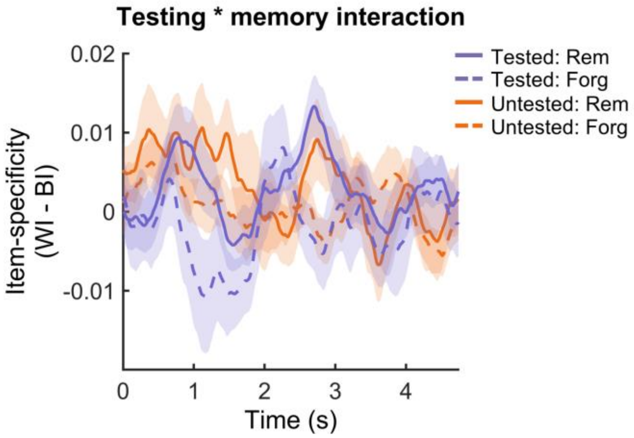


**Fig S7. Item-specific representations were not modulated by pre-sleep testing.** To understand whether the pre-sleep testing influence memory reactivations, we performed a three-way repeated measures ANOVA, with pre-sleep testing (tested vs. untested), item-specificity (WI vs. BI), and post-sleep memory (remember vs. forget) as factors. No significant interaction effect was found in any individual time window ($p > 0.081$). Furthermore, a two-way repeated measures ANOVA with pre-sleep testing and item-specific representations as factors for post-sleep remembered items did not reveal any significant interaction effect ($p_{corr} > 0.383$).

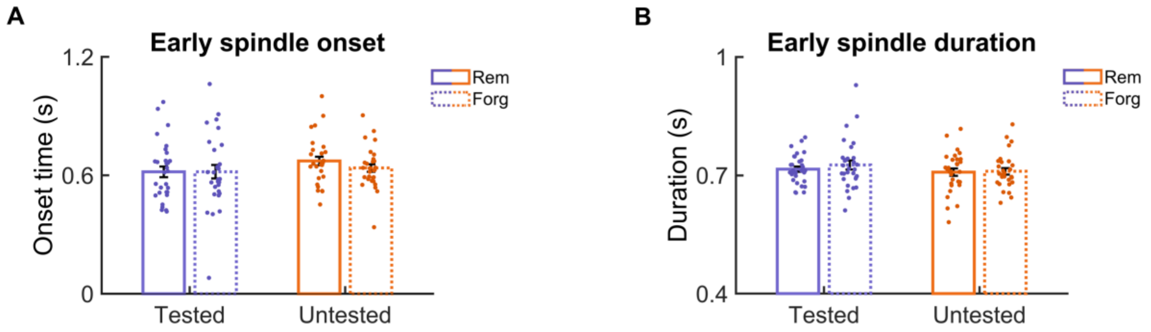


**Fig S8. The onset time and duration of the spindles that occurred within the first 2 s following TMR cues were not different either between tested and untested items or between post-sleep remembered and forgotten items.** We extracted the spindle onset time and duration for the spindles that occurred within the first 2s and then performed testing (tested vs. untested) by subsequent memory (remember vs. forget) repeated measures ANOVA on time and duration, respectively. The results found that neither a significant interaction for the spindle onset time ($F(1, 29) = 0.71$, $p = 0.408$, **A**) nor a significant interaction effect for the spindle duration ($F(1, 29) = 0.83$, $p = 0.371$, **B**).

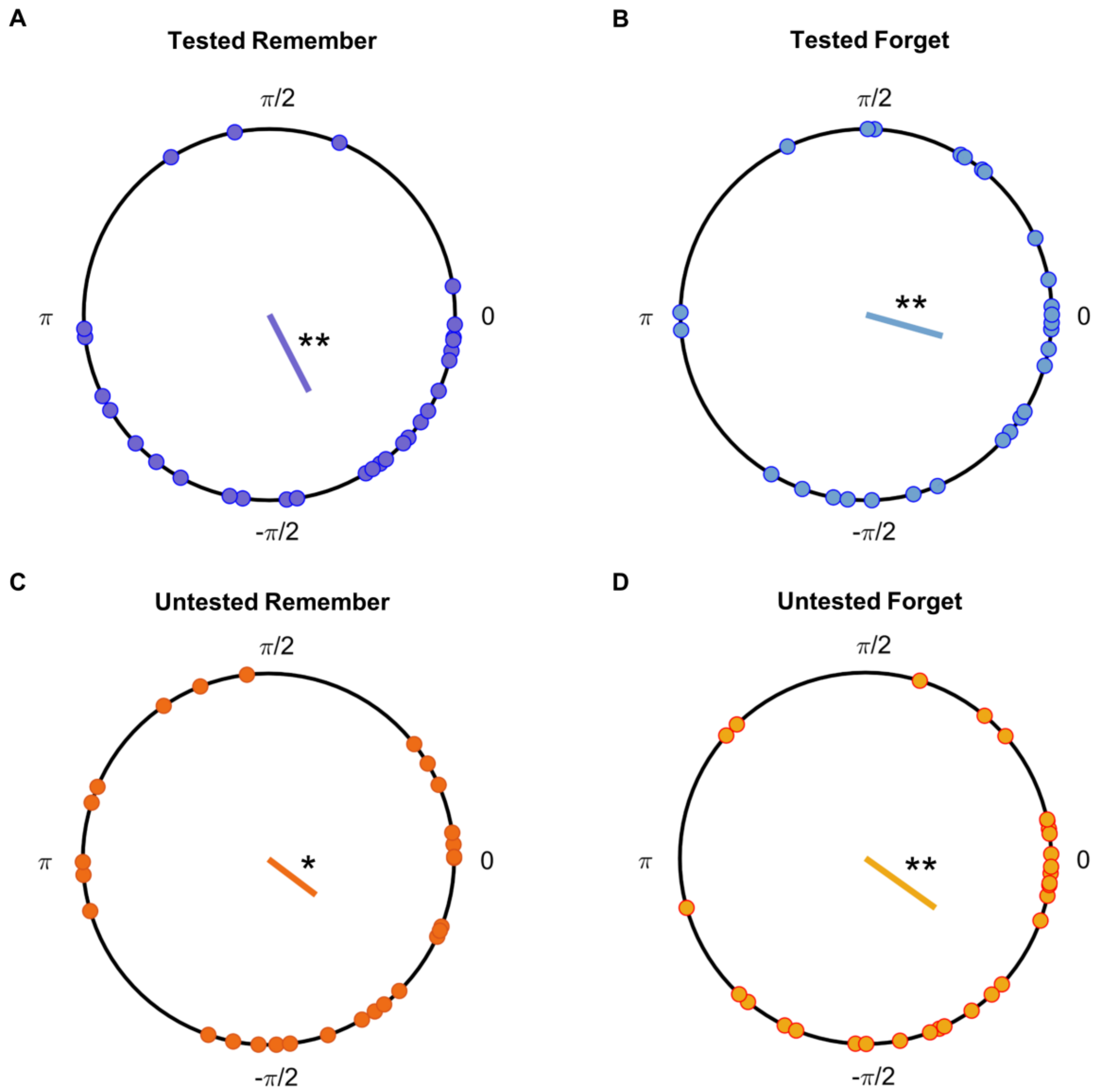


**Fig S9. Spindles in an extended late time window (2.2-4.2 s) were coupled to the up-state of slow oscillations.** For trials that showed both the spindles and slow oscillations in the late time window (i.e., 2.2-4.2 s), we extracted the preferred phase of the slow oscillation for each spindle in this time window and calculated the mean phase across trials for each participant (CircStat toolbox). The Rayleigh Z test was then used to determine if the distribution of phases deviated from a uniform distribution across participants. The results showed that spindle activities for both subsequently remembered and forgotten tested items **(A-B)** as well as for subsequently remembered and forgotten untested items **(C-D)** were all significantly and preferentially coupled to the up-state of SOs (all *ps* $< 0.045$). *: $p < .05$; **: $p < .01$.

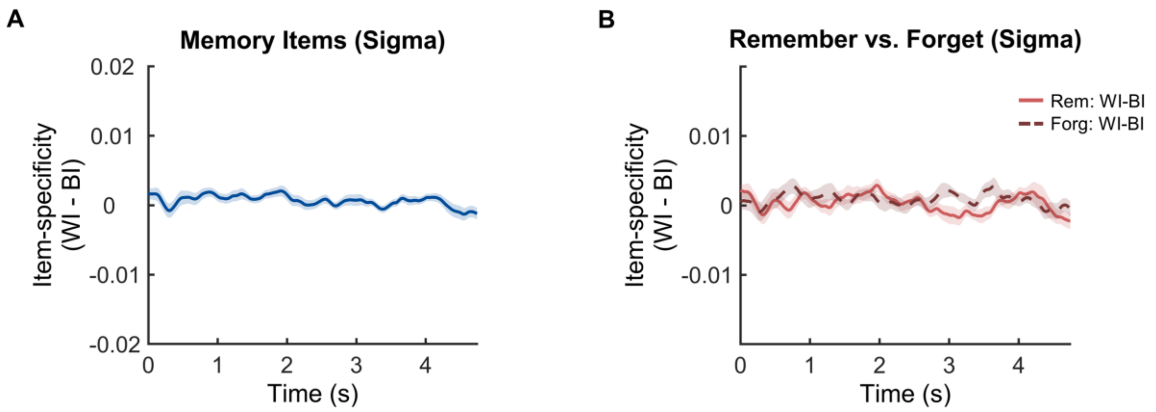


**Fig S10. No significant item-specific representations when using the sigma band power in the RSA.** We performed the same representational similarity analysis using the sigma band power that was extracted from the 11-18 Hz of the raw EEG data. **(A)** No significant item-specific representations were identified for memory items ($p_{corr} > 0.063$). **(B)** No significant clusters showed a significant difference between item-specific representations for post-sleep remembered versus forgotten items ($p_{corr} > 0.111$).
